# Simple recombinant monoclonal antibody production from *E. coli*

**DOI:** 10.1101/2024.04.17.589891

**Authors:** Karen Baker, Tara A. Eastwood, Esther Garcia, Christopher Lennon, Daniel P. Mulvihill

## Abstract

Antibodies are valuable biological reagents used in a wide range of discovery research, biotechnology, diagnostic and therapeutic applications. Currently both commercial and lab scale antibody production is reliant on expression from mammalian cells, which can be time consuming and requires use of specialist facilities and costly growth reagents. Here we describe a simple, rapid and cheap method for producing and isolating functional monoclonal antibodies and antibody fragments from bacterial cells that can be used in a range of laboratory applications. This simple method only requires access to basic microbial cell culture and molecular biology equipment, making scalable in-house antibody production accessible to the global diagnostics, therapeutics and molecular bioscience research communities.

## Introduction

Antibodies (or Immunoglobulins) are fundamental reagents in the diagnosis and treatment of a wide range of common and rare health disorders, and are valuable tools within the discovery research laboratory. Antibodies exist in diverse forms, ranging from monomeric single chain polypeptides (e.g. single domain antibodies / nanobodies) to multimeric complexes with heavy and light chain components. While their structure and function may vary significantly, they each have common physical properties that can provide challenges during recombinant expression. These include precise folding with controlled sequential inter- and intra-chain disulphide bond formation, as well as post-translational modifications, such as glycosylation, to ensure stability and function within the body.

Current biotherapeutic monoclonal antibody (mAb) production relies primarily on the use of mammalian expression systems, such as Chinese hamster ovary (CHO) cells, which facilitate the protein folding and complex post-translational modifications required for production of functional antibody complexes (Walsh, 2018). In contrast, due to their simpler structure and smaller size, antigen-binding fragments (Fab) and other simpler IgG fragment containing fusions, also used in biotherapeutics and research applications, can be expressed in the Gram-negative bacteria, *Escherichia coli*. These bacterial cells are simpler and quicker to grow than mammalian systems and allow yields at the g/L scale in fermentation batch cultures (Gupta and Shukla, 2017). Antibody fragments have been produced from *E. coli* through targeting the Fab peptide to the periplasmic region to allow formation of the required disulphide bonds required for functional antibody-complex. (Kaplon et al., 2022; Sandomenico et al., 2020; Walsh, 2018) Similar approaches have now been successfully applied to produce a limited number of full-length IgG antibodies from bacteria (Simmons et al., 2002), and while they lack glycosylation required for preventing some immuno-recognition activities, they remain stable in the blood stream for periods equivalent to CHO derived molecules (Simmons et al., 2002). Unless expressed in specialised commercial strains, *E. coli* produced IgGs and Fabs are normally reduced and insoluble, and require sequential rounds of controlled *in vitro* folding and regulated oxidation steps for the correct disulphide bonds to form in order to produce a functional IgG complex.

Here we describe a simple method to produce functional Fab and mAb complexes from *E. coli*. Using a peptide tagging-based method developed in this lab for producing recombinant protein-containing vesicles (Eastwood et al., 2023), functional heterodimeric Fabs and mAbs can be produced and purified from compartmentalised cytosolic vesicles within *E. coli*. We have applied the technique to isolate a model Fab and several monoclonal antibodies, and demonstrate their functionality using a combination of ELISA, western blot, immunoprecipitation and immunofluorescence assays. This simple, quick and cheap antibody production method can be applied to produce monoclonal antibodies in laboratories with basic microbial culturing facilities, opening the potential for in-lab antibody production for the entire bioscience research community.

## Materials and Methods

### *E. coli* cell culture and protein induction

All bacterial cells were cultured using LB (10 g Tryptone; 10 g NaCl; 5 g Yeast Extract (per litre)) and TB (12 g Tryptone; 24 g Yeast Extract; 4 ml 10% glycerol; 17 mM KH_2_PO_4_ 72 mM K_2_HPO_4_ (per litre) media. 5 ml LB starters from fresh bacterial transformations were cultured at 37ºC to saturation and used to inoculate 25 - 500 ml volume TB media flask cultures that were incubated overnight at 30ºC with 200 rpm orbital shaking (2.5 cm throw). Oxygenation of cultures was enhanced by ensuring a large liquid surface area during growth (i.e. 25 ml media in 500 ml conical flask or 500ml / 1 L of media in a 5 L conical flask). Recombinant protein expression from the T7 promoter was induced by addition of IPTG to a final concentration of 20 μg/ml once the culture had reached an OD_600_ of 0.8 – 1.0.

### Fission yeast cell culture

Prototrophic *Schizosaccharomyces pombe cam1*.*gfp* cells (Baker et al., 2019) were cultured and maintained according to (Moreno et al., 1991) using Edinburgh minimal medium with glutamic acid nitrogen source (EMMG). Cells were cultured at 25ºC and maintained as early to mid-log phase cultures for 48 hr prior to formaldehyde fixation.

### Human cell culture

The A2780 human ovarian carcinoma and RPE-1 human cell lines were previously purchased from the Health Protection Agency (Salisbury, UK). Both cell lines were cultured in Iscove’s Modified Dulbecco’s Medium (IMDM) supplemented with 10% Fetal bovine serum (FBS) at 37 °C and 5% CO2. Cells cultured for subsequent fixation and microscopy analysis were plated at 5 x 10^5^ cells per 35 mm imaging dish (μ-Dish 35 mm, Thistle Scientific, UK) and cultured for 24 hr prior to fixation.

### Soluble protein extracts

*E. coli* cell pellets from 50 ml cultures were resuspended in 5 ml of 100 mM Tris (pH 8.0), sonicated for a total of 2 min (6 x 20 sec pulses), and cell debris removed by centrifugation at 7,700 RCF (4ºC) for 20 min. For human cells, pellets of A2780 or RPE-1 cells were resuspended in lysis buffer (50 mM HEPES, 250 mM NaCl, 0.1% NP40, 1 mM AEBSF, 10 μg/ml Aprotinin, 20 μM Leupeptin, 1 mM DTT, 1 mM EDTA, 1 mM NAF, 10 mM Na β-glycerophosphate, 0.1 mM Na orthovanadate pH 7.4) and incubated on ice for 30 min. Cell debris was pelleted at 4,000 RCF for 10 min at 4ºC.

### Protein purification

Fab & mAb heterodimers were isolated either by sequential Nickel and protein-G affinity purifications (Figure 1E) or using a single-step purification on protein G (Figure 2G), where cleared soluble antibody-containing bacterial cell extracts were passed through a 0.45 μm filter and bound to protein-G matrix (agarose or Dynabeads). The matrix was subsequently washed 3 x in excess 100 mM Tris (pH 8.0) and 3 x in excess 10 mM Tris (pH 8.0) before antibodies were eluted from protein G with 50 mM glycine (pH 3.0) into a 1/5 volume of 1 M Tris (pH 8.0) neutralising buffer. Yields of purified functional IgG complex were determined using both gel densitometry and spectrometry.

**Figure 1.**
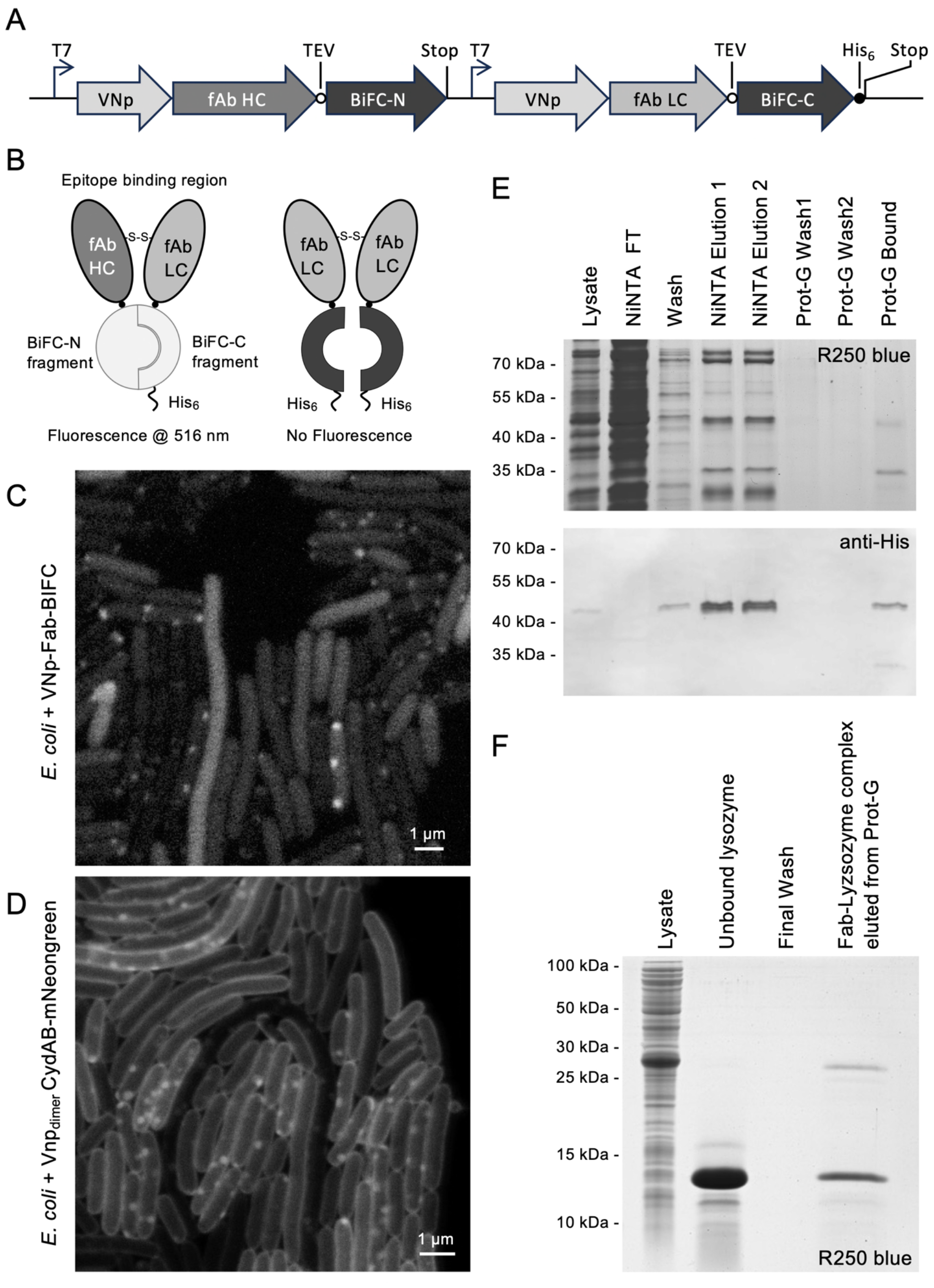
Functional VNp-Fab chain fusion heterodimer isolation from *E. coli*. (A) Schematic of plasmid DNA region that encodes for VNp-anti-lysozyme-Fab–BiFC heavy and light chain fusions. (B) Schematic highlighting how Fab heterodimers result in BiFC dependent mVenus fluorescence. (C) STED image of live *E. coli* cells expressing VNp-Fab–BiFC fusion proteins displaying BiFC-mVenus fluorescence highlighting presence of Fab heterodimers within cytosolic vesicles (Scale: 1 μm). (D) STED image of live *E. coli* cells expressing dimerising VNp and mNeongreen labelled inner membranes protein CydAB highlights presence of cytosolic inner membrane vesicles (Scale: 1 μm). (E) Coomassie R250 SDS-PAGE analysis (upper panel) and anti-His_6_ western blot illustrating Nickel affinity purification and subsequent protein G binding functionality of VNp-Fab–BiFC heavy and light chains. Protein-G beads were subsequently washed in binding buffer, before being boiled in SDS-PAGE loading buffer to release bound proteins. (F) Coomassie SDS-PAGE analysis of lysate of *E. coli* expressing VNp-Fab heavy and light chains, which were purified directly onto Protein-G agarose and subsequently bound to soluble chicken egg lysozyme, the target of the Fab heterodimer. The Protein-G – Fab – lysozyme immunocomplex was washed in excess IP buffer before being boiled in SDS-PAGE loading buffer.

**Figure 2.**
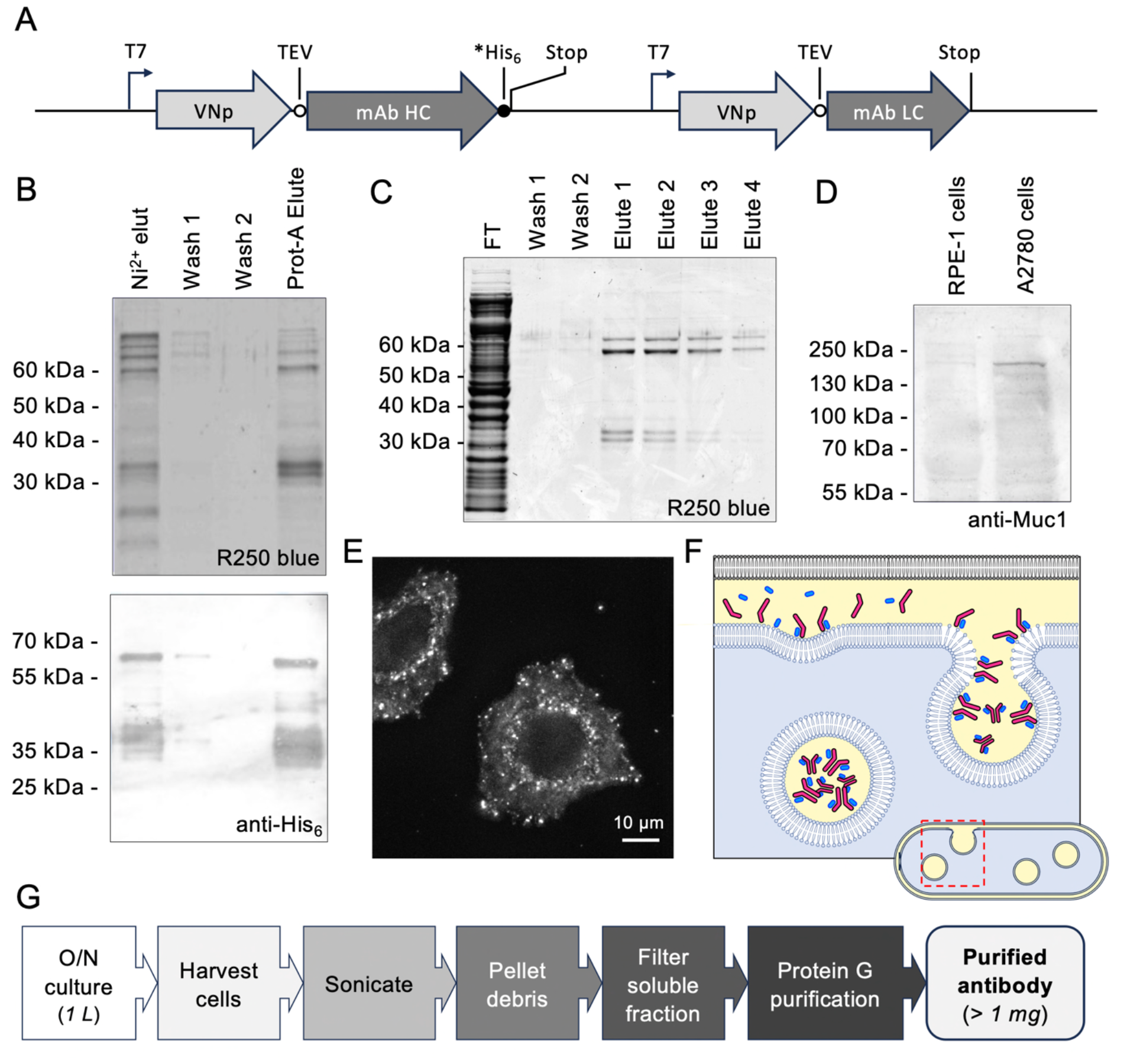
Bacterially expressed VNp-mAb chain fusions form functional anti-Muc-1 antibodies. (A) Schematic of plasmid DNA region encoding for VNp-anti-Muc1-mAb heavy and light chains. (B) Coomassie R250 SDS-PAGE analysis (upper panel) and anti-His6 western blot illustrating Nickel affinity purification and subsequent protein G binding functionality of of VNp-anti-Muc1-mAb heavy and light chains. Protein-G beads were subsequently washed in binding buffer, before being boiled in SDS-PAGE loading buffer to release bound proteins. (C) Coomassie SDS-PAGE analysis of fractions from Protein-G purification of VNp-anti-Muc1-mAb from *E. coli* extract. (D) VNp-anti-Muc1 western blot analysis of extracts from cultures of RPE-1 and A2780 ovarian cells (Muc1 predicted size: 122.1 kDa). (E) VNp-anti-Muc1 immunofluorescence of A28780 cells reveals Muc1 foci at membranes (Scale: 10 μm). (F) Model illustrating how VNp dependent inner membrane interactions could spatially and temporally coordinate mAb heavy (magenta) and light (blue) chain tetramer complex formation and incorporation into periplasm (yellow) filled cytosolic vesicles. (G) Summary of simplified, rapid bacterial mAB expression and purification protocol.

### Immuno-assays

Western blots and immunoprecipitations were undertaken as described previously, using alkaline phosphatase conjugated secondary antibodies for detection (Harlow and Lane, 1988). Anti-GFP immunofluorescence was undertaken in fission yeast using the method described in (Hagan and Hyams, 1988) with the omission of glutaraldehyde from the fixation step. Anti-Muc1 immunofluorescence was undertaken on cultured A2780 human ovarian cancer cells cultured on coverslips as described above. Cells were fixed with 4% formaldehyde solution for 15 min. After fixation formaldehyde was removed and cells washed 3 x with PBS and permeabilised in TBS + 0.3% Triton-X for 15 min. Permeabilised cells were washed twice in TBS before blocking in 2% BSA in TBS-Tween (0.1%) for 1 hr. Cells were incubated in VNp-anti-Muc1 primary mAb in 2% BSA in TBS-Tween (0.1%) overnight at 4°C. Cells were washed 3 x in TBS-Tween (0.1%) + 2% BSA before incubation in TRITC conjugated anti-Mouse secondary antibody (Sigma) at 1:1000 concentration for 1 hr in the dark. Cells were subsequently washed 2 x in 2% BSA in TBS-Tween (0.1%) and 1x in TBS prior to visualisation. All primary antibodies were used at 1 in 20 dilution for IF experiments. Secondary only controls confirmed observed immunofluorescence signal was due to presence of primary antibody.

### Enzyme-Linked ImmunoSorbent Assay (ELISA)

*E. coli* expressed and purified VNp-anti-GFP mAb (as described above) were tested by alkaline phosphatase ELISA. GFP (0 to 1.5 μM) was bound to black 96 well plates, washed 3x with TBS and blocked overnight at 4 ºC with 0.5% IgG free BSA and 0.1 % Tween. Plate was subsequently incubated with the test mAb (e.g. VNp-anti-GFP) at 4 ºC overnight in TBS + 0.1 % Tween (TBST); washed 5 x with TBST; incubated for 1 hr with a 1:10,000 dilution alkaline phosphatase conjugated anti-mouse secondary antibody (Sigma), prior to a final 5 x TBST washes and incubation with BCIP/NBT substrate. Signal at 620 nm was measured using a BMG labtech Clariostar plate reader.

### Microscopy

Samples were visualised using an Olympus IX71 microscope with UApo N 100x 1.49 oil immersion lens mounted on a PIFOC z-axis focus drive (Physik Instrumente, Karlsruhe, Germany), and illuminated using LED light sources (Cairn Research Ltd, Faversham, UK) with appropriate filters (Chroma, Bellows Falls, VT). Samples were visualised using a QuantEM (Photometrics) EMCCD camera, and the system was controlled with MetaMorph software (Molecular Devices). Each 3D-maximum projection of volume data was calculated from 21 z-plane images, each 0.2 μm apart. STED images were acquired using a Leica TCS SP8 STED 3X microscope equipped with a pulsed White Light Laser at 80MHz and fitted with a STED white HC PL APO 86x/1.2 water immersion lens. mVenus and mNeongreen fluorescent proteins were depleted using a 592nm continuous-wave STED laser.

### Molecular Biology

Tags and gene sequences were codon optimised for expression in *E. coli*. Bacterial expression plasmids (and sequences) generated and described in this study (pRSFDUET-VNp-anti-Lysozyme Fab-BiFC; pRSFDUET-VNp-anti-Lysozyme Fab; pRSFDUET-VNp-anti-Muc1 mAb; pRSFDUET-VNp-anti-6xHis mAb; pRSFDUET-VNp-anti-GFP mAb) will be made available at https://www.addgene.org/Dan_Mulvihill/ upon publication.

## Results

We previously described the use of a short vesicle nucleating peptide (VNp) tagging technology to facilitate the enhanced expression of functional recombinant proteins from *E. coli*. (Eastwood et al., 2023; Streather et al., 2023). As well as enhancing nanobody production, the VNp technology allows production of functional Etanercept, a disulphide bond-dependent homodimeric IgG1 fusion anti-inflammatory therapeutic, which can be isolated from cytosolic membrane bound vesicles within the *E. coli* cell (Eastwood et al., 2023). Spurred on by this initial success, we decided to explore whether the technology could be applied to produce valuable, heterodimeric immunoglobulin complexes from *E. coli*, and thus negate the need for periplasmic targeting, re-solubilisation and chemically induced disulphide formation steps, or expression in mammalian cell systems. To test this, a plasmid was generated to facilitate bacterial co-expression of VNp fusions of heavy and light chain Fab peptides derived from the anti-lysozyme monoclonal antibody, D1.3 (Amit et al., 1986). cDNAs encoding for each Fab fragment chain were fused to 5’ VNp and 3’ Bimolecular Fluorescence complementation (BiFC) encoding sequences (Kodama and Hu, 2010) (Fig 1A) to confirm successful formation of light-chain/heavy-chain heterodimer (Fig 1B). Expression from this construct in *E. coli* resulted in BiFC-Venus fluorescence from cytosolic vesicular structures (Fig 1C), consistent in appearance with internalised VNp dependent membrane bound vesicles (Fig 1D), and reminiscent of cytosolic *E. coli* membrane vesicles containing the homodimeric IgG1 therapeutic fusion protein, VNp-Etanercept (Eastwood et al., 2023). BiFC fluorescence (Fig 1C) and SDS-PAGE analysis (Fig S1) of cell lysate, nickel affinity purification (via light chain carboxy-terminal His_6_-tag) and subsequent protein G conjugations were consistent with the successful production and purification of correctly folded heterodimeric Fab (Fig 1E). Functionality of the same Fab complex lacking the BiFC fusion tags was confirmed by establishing its ability to associate with soluble chicken egg Lysozyme, the epitope target of D1.3, during immunoprecipitation assays (Fig 1F). These data confirmed fusing the Vesicle Nucleating peptide to the amino terminal of Fab heavy and light chains allows production of functional Fab heterodimer complex into membrane vesicles within the *E. coli* cytosol.

Encouraged by the ability to produce functional heterodimeric antibody fragments, we explored whether the VNp tag could be used to facilitate bacterial expression of tetrameric monoclonal antibodies (mAb). We thus generated *E. coli* optimised genes encoding for VNp fusions of a mAb against Muc1 (Fig 2A) (Thie et al., 2011), a type I mucin that is mis-regulated in a variety of cancers, and is thus an attractive diagnostic biomarker (Nath and Mukherjee, 2014). Both VNp-mAb chains expressed as soluble proteins that co-purified, via a carboxyl His_6_-tag on the mAb heavy chain, as a complex on nickel resin (Fig 2B). The heterodimeric nature and functionality of the purified mAb complex was subsequently confirmed by its ability to bind Protein A (Fig 2B). A simplified purification strategy by which functional recombinantly expressed mAb chains are directly isolated from filtered bacterial extracts onto Protein-G matrix was developed (Fig 2C) and employed during subsequent antibody preparations.

Western blots analysis of extracts from A2780 ovarian cancer cells and parental RPE-1 cells using the VNp-anti-Muc1 as primary antibody revealed a discrete band which migrated at a size consistent with Muc-1 protein (Fig 2D). The VNp-anti-Muc-1 band was significantly more intense in the extract from A2780, which is consistent with the fact that Muc-1 is a biomarker upregulated in ovarian cancer cells (Horm and Schroeder, 2013). Additional functionality and applicability of the bacterial expressed anti-Muc1 mAb was confirmed by immunofluorescence analysis of the A2780 cells (Fig 2E), which revealed a localisation pattern consistent with that observed previously using commercial mammalian cell derived antibodies (Bennett et al., 2001; Fanayan et al., 2009; Wang et al., 2011).

To further confirm the applicability of the VNp technology for antibody production in bacteria, we tested the same purification strategy (Fig 2G) with two further mAbs, N86/38.1R anti-GFP and N144/14R anti-6xHis (Andrews et al., 2019), which are commonly applied applicable in a range of routine cell and molecular research applications. Using the published mAb peptide sequences, *E. coli* codon optimised cDNA was generated to facilitate expression of VNp-tagged heavy and light chains of anti-GFP and anti-6xHis mAbs (Fig 3A). Tetrameric-disulphide-bond stabilised complexes of VNp-anti-GFP mAb heavy and light chains were isolated from filtered *E. coli* cell extracts with protein-G matrix (Fig 3B, S1), with both ELISA and western blot analyses confirming functionality of the purified mAb by its ability to specifically and quantifiably recognise purified GFP fusions, as well as GFP-tagged proteins within crude cell extracts (Fig 3C, S1). Using the same strategy N144/14R anti-6xHis mAb (Andrews et al., 2019) was successfully produced as a VNp-fusion complex, and functionality confirmed by its use as a primary antibody during western blot analysis of extracts and purified proteins containing His_6_ affinity tags (Figure 3D). Milligram yields of these protein-G purified mAbs were typically obtained from 1 litre shaking flask cultures of *E. coli*. Purified mAb was stored at a 2 μM stock concentration (in Tris buffer pH 8.0 at 4ºC), and typically diluted 5,000 fold for use as a primary antibody in western blot analysis (e.g. Fig 3C-D). To confirm applicability of the VNp-anti-GFP mAb as a reagent for *in situ* studies, it was used as a primary antibody during indirect immunofluorescence analysis of fission yeast cells in which the endogenous essential calmodulin, Cam1, had been labelled with GFP. Fluorescence signals from the GFP fluorophore and TRITC labelled immunocomplex overlapped precisely, confirming the efficacy of the VNp-mAb for in cell studies (Fig 3E).

**Figure 3.**
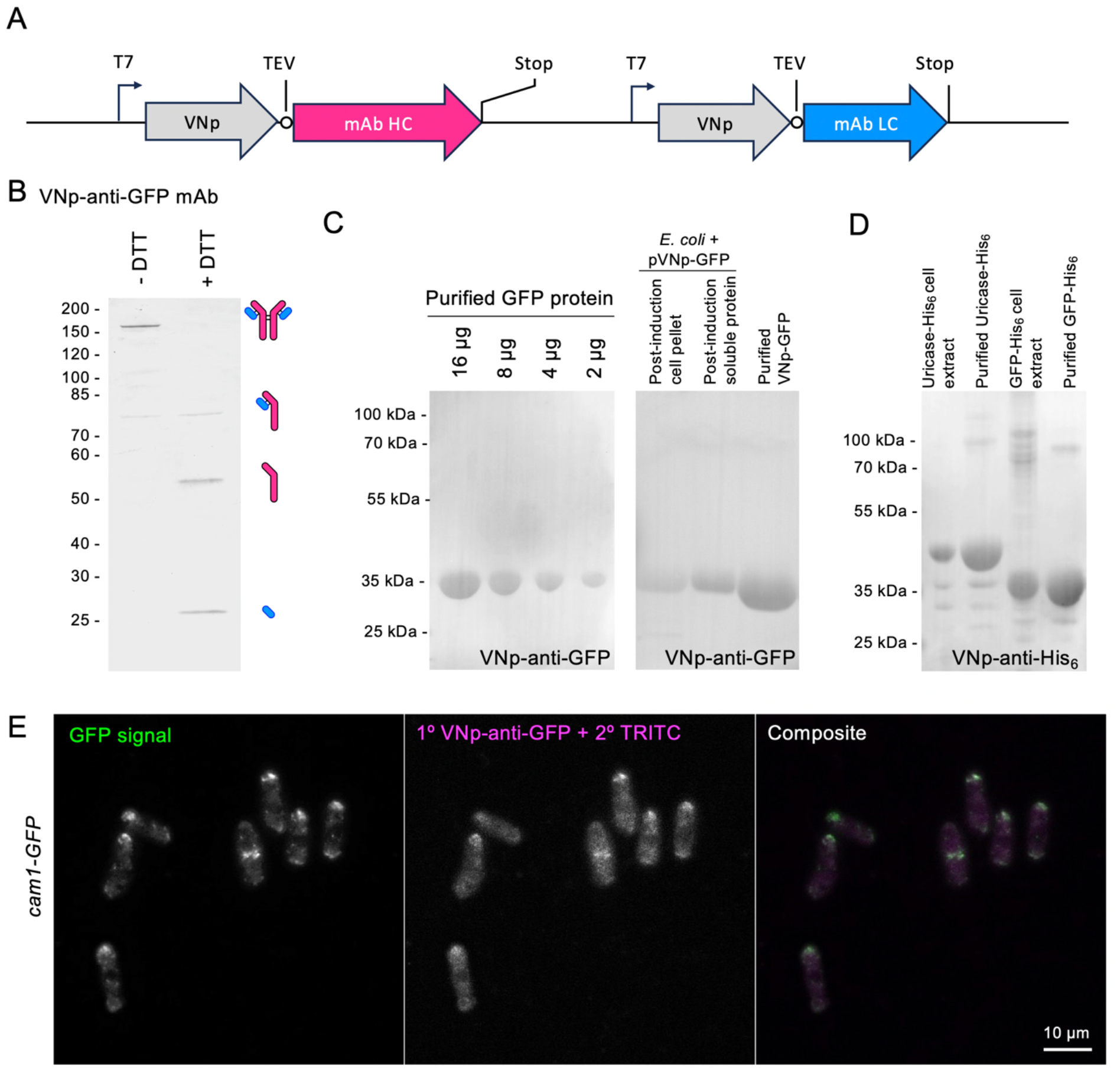
Simplified expression and one step purification of functional VNp-anti-GFP and VNp-anti-6xHis monoclonal antibody complexes from bacteria. (A) Schematic of plasmid region encoding for VNp-mAb heavy (magenta) and light (blue) chains. (B) Coomassie SDS-PAGE analysis of Protein-G purified VNp-anti-GFP mAb illustrate heavy and light chains migrate at the expected size of a tetrameric complex in the absence of the reducing agent, DTT. (C) VNp-anti-GFP western blots of affinity purified GFP protein and crude cell extracts of cells expressing GFP fusions. (D) VNp-anti-His_6_ western blots of His_6_-tagged proteins in purified samples or crude *E. coli* cell extracts. (E) Indirect GFP immunofluorescence staining of *cam1-gfp* fission yeast cells using VNp-anti-GFP mAb primary and TRITC-conjugated secondary antibodies reveals colocalization between signals from the GFP tag (green) and TRITC-labelled antibody (magenta) (Scale: 10 μm).

## Discussion

Here we have described the application of a *vesicle nucleating peptide* tagging methodology (Eastwood et al., 2023) to facilitate simple and rapid expression and purification of diverse functional IgG complexes from bacterial cells. Using this method, milligram scale yields of affinity purified functional mAb were isolated from each litre of overnight *E. coli* culture. The subsequent purified mAb can be diluted to sub-nanomolar concentrations when employed as a primary antibody in western blot analyses with alkaline phosphatase detection (Fig 2 & 3). The method is extremely rapid, taking only 3 days from bacterial transformation to isolation of purified active antibody onto matrices of IgG binding ligand, such as Protein G. This strategy has resulted in successful production of each heterodimeric IgG complex examined. Unlike previous IgG *E. coli* expression methods, this simple technique can be rapidly applied to the production of diverse heteromeric immunoglobulin complexes from *E. coli*, without the need for expression level and periplasmic targeting optimisation, protein re-solubilisation or chemically induced disulphide formation steps.

How the VNp targeting system facilitates bacterial IgG complex production is unclear. Current dogma suggests that functional IgG complex formation requires controlled sequential folding and disulphide bond formation events to occur, with some steps taking place in specific subcellular locations, such as the ER (Feige et al., 2010). Our findings support previous work which indicated that the IgG complex can form in the absence of ER and folding control checkpoints (Simmons et al., 2002). One hypothesis is that the VNp driven interaction with the bacterial membrane not only brings the individual antibody chains together in an oxidising environment (i.e. the periplasmic sourced lumen of the internalised vesicles), but also brings the individual chains together in a spatially, temporally and stoichiometrically controlled manner that promotes both correct peptide folding and inter-chain interactions (Fig 2F). These findings suggest that IgG complex assembly and folding is more intrinsic to the protein components themselves than had previously been imagined.

Bacterial produced mAbs can be applied to discovery research, diagnostic and therapeutic applications just like the mammalian cell sourced equivalents. While the bacterial produced mAbs lack the glycosylation seen in mammalian produced proteins, glycosylated and non-glycosylated IgGs have equivalent *in vitro* binding and *in vivo* lifetimes in the mammalian blood stream (Hobbs et al., 1992; Tao and Morrison, 1989).

This bacterial antibody production method provides several benefits over mammalian cell-based systems. Not only does it represent significant cost savings over mammalian cell culture, bacterial cells are quicker and simpler to grow, and do not require specialised tissue culturing facilities. Also, the simple one step purification strategy described here (Fig 2G) does away with the need for specialised expensive chromatography purification equipment and reagents. Thus, this VNp technology makes monoclonal antibody production a reality for any research lab with access to only basic molecular and microbiology facilities, therefore making monoclonal antibody production methodology accessible to the global research community.

## Acknowledgements

The authors thank Cassie Stone and Professor Michelle Garrett for providing cell culture samples. We thank Professors B. Goult, C. Gourlay, P. Gunning and I. Hagan for helpful comments on the manuscript. This work was supported by the University of Kent and funding from the Biotechnology and Biological Sciences Research Council (BB/S005544/1 & BB/X007448/1), the Medical Research Council (MR/X502753/1) and Fujifilm-Diosynth Biotechnologies UK Ltd. The Vesicle Nucleating peptide technology described here and its use in antibody production is associated with patent applications *GB2118435*.*3* and *PCT/GB2022/053239*.

## Author Contributions

KB and TAE performed the experimental studies; KB, TAE, CL and DPM designed experiments; EG supervised STED experiments and processing; DPM sought funding and supervised the project. DPM wrote the main drafts of the manuscript and all authors contributed to editing.

## Figure Legends

**Figure S1.**
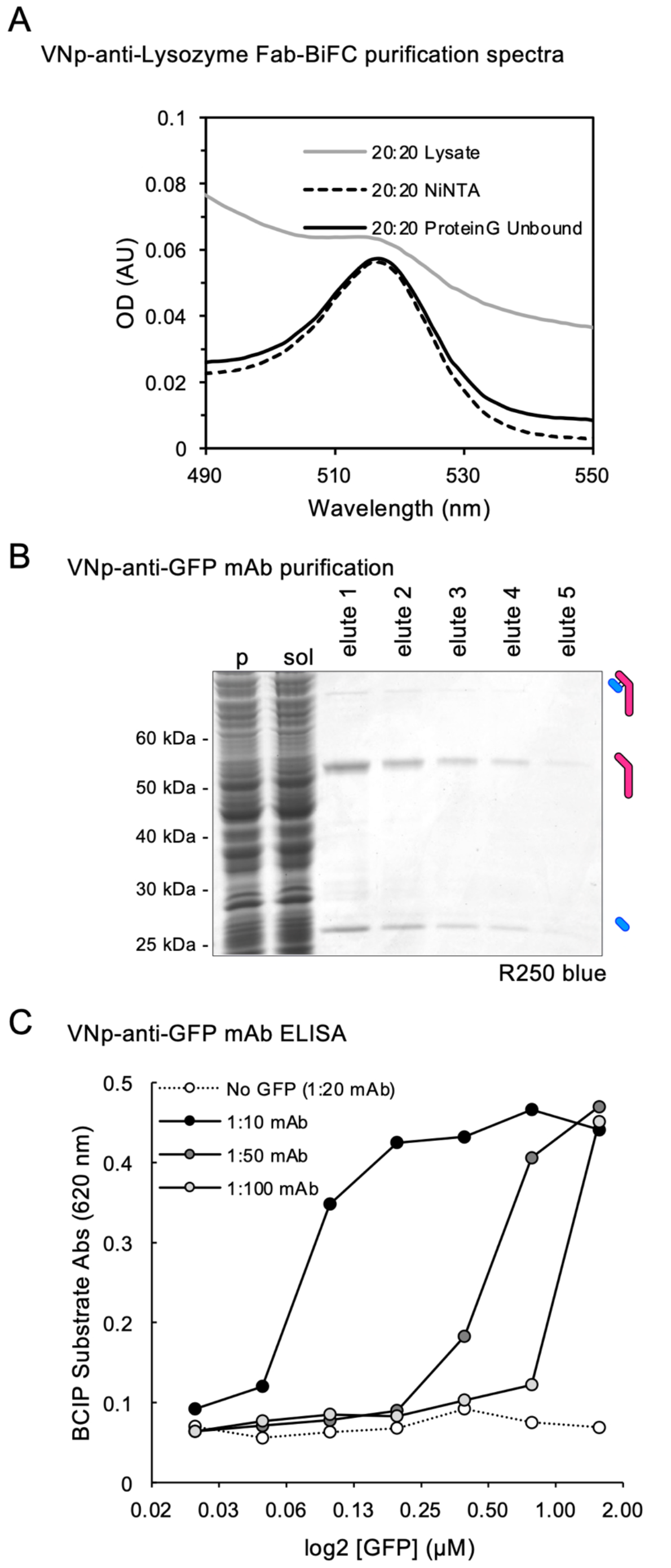
VNp-Fab-BIFC purification fluorescence and VNp-anti-GFP mAb purification and ELISA. (A) Fluorescence spectra highlight presence of Bimolecular Fluorescence Complementation dependent mVenus signal in nickel affinity purified (dashed line) and subsequent Protein G purified (black line) proteins from lysate (grey line) of cells shown in Figure 1C. (B) Coomassie SDS-PAGE analysis of Protein-G purification fractions from an extract *E. coli* expressing VNp-anti-GFP mAb heavy and light chains. (C) ELISA of *E. coli* expressed VNp-anti-GFP mAb dilutions against purified GFP protein and detected with alkaline-phosphatase conjugated anti-mouse secondary antibody.

